# vamos: VNTR annotation using efficient motif sets

**DOI:** 10.1101/2022.10.07.511371

**Authors:** Jingwen Ren, Bida Gu, Mark JP Chaisson

**Author notes:** These authors contributed equally to this work (Jingwen Ren, Bida Gu).

## Abstract

**Motivation:** Roughly 3% of the human genome is composed of variable-number tandem repeats (VNTRs): tandemly repeated arrays of motifs at least six bases. These loci are highly polymorphic: over 61% of insertion and deletion variants at least 50 bases found from long-read assemblies are inside VNTRs. Furthermore, long-read assemblies reveal that VNTR loci are multiallelic, and can vary by both motif composition and copy number. Current approaches that define and merge variants based on alignment breakpoints do not capture this complexity of variation. A natural alternative approach is to instead define the motif composition of VNTR sequences from samples, and to detect differences based on comparisons of repeat composition. However, due to the complexity of VNTR sequences, it is difficult to establish a common reference set of motif sequences that may be used to describe variation in large sequencing studies.

**Results:** Here we present a method vamos: <u>V</u>NTR <u>A</u>nnotation using efficient <u>Mo</u>tif <u>S</u>ets that for any VNTR locus selects a set of representative motifs from all motifs observed at that locus that may be used to encode VNTR sequences within a bounded edit distance of the original sequence. We use our method to characterize VNTR variation in 32 haplotype-resolved human genomes. In contrast to current studies that merge multi-allelic calls, we estimate an average of 3.1-4.0 alleles per locus.

**Availability:** github.com/chaissonlab/vamos, zenodo.org/record/7158427

## 1 Introduction

Variable number-tandem repeats (VNTRs) are a class of repetitive DNA composed of short DNA sequences called motifs repeated many times in tandem. By convention, VNTRs are composed of motifs at least six bases; shorter motifs are classified as short tandem repeats. The repetitive nature of these sequences primes them for mutations through strand slippage and unequal crossing over that increase or decrease motif copy number or introduce mutations of motifs (Levinson and Gutman, 1987). Variation of VNTR sequences has been found to impact physiology and cellular function. Disease studies have found associations of VNTR length or composition with diabetes (Torsvik *et al*., 2010), schizophrenia(Song *et al*., 2018), and Alzheimer’s (De Roeck *et al*., 2018). Additionally, methods developed to analyze VNTR variation using high-throughput short read sequencing data found widespread association between VNTR length and gene expression (Bakhtiari *et al*., 2021; Lu *et al*., 2021; Garg *et al*., 2021). Finally, variation directly in coding sequences are found to be linked with human traits including height and hair patterns (Mukamel *et al*., 2021; Beyter *et al*., 2021).

Despite the growing research on the functional impact of VNTR variation, the overall knowledge of genetic diversity in these sequences lags behind non-repetitive DNA. VNTR sequences have been difficult to study because the majority of sequencing has been with short reads that have degenerate alignments and reference bias. As a consequence, they are masked in short-read studies using low complexity filters (Auton *et al*., 2015; Zook *et al*., 2019). Contrary to short-read sequencing, variation of VNTR loci is routinely resolved using long-read alignments (Sedlazeck *et al*., 2018) and assemblies (Chaisson *et al*., 2019; Ebert *et al*., 2021). The initial long-read assembly studies found that insertions and deletions greater than 50 bases (structural variation) in humans are enriched in VNTR loci (Pendleton *et al*., 2015; Chaisson *et al*., 2019); over 61% of structural variants discovered in 32 haplotype-resolved assemblies produced by the Human Genome Structural Variation Consortium (Ebert *et al*., 2021) are inside VNTRs. The loci that harbor these structural variants are highly polymorphic. An analysis of ∼30,000 VNTRs in these assemblies (corresponding to loci studied in (Lu *et al*., 2021)) shows ∼9 alleles per locus, when considering each distinct sequence as an allele. This analysis with recent long-read data in human agrees with the long-studied observation that VNTR mutation rate in prokaryotes is up to six orders of magnitude greater than single-nucleotide polymorphisms.(Vogler *et al*., 2006; Jee *et al*., 2016)

As larger studies of long-read sequencing become available, the number of different alleles will likely rise, presenting a challenge for how to represent VNTR variation in populations, and even how to define what constitutes a different allele. Current methods to discover variants from long-read data or assemblies detect insertion or deletion breakpoints based on sequence alignment (Jiang *et al*., 2020; Sedlazeck *et al*., 2018; Ebert *et al*., 2021). Analysis that is more advanced than breakpoint resolution has enabled association of VNTR variation and traits or disease. A study of 3,622 Icelandic individuals sequenced with long-reads used a combination of pre-filtering and clique-based clustering to identify structural variant alleles in VNTR sequences that associate with height, atrial fibrillation, and recombination by VNTR length (Beyter *et al*., 2021). In addition to this study, others focused on eQTL analysis of VNTR loci(Bakhtiari *et al*., 2021; Lu *et al*., 2021; Garg *et al*., 2021), the associations are found between the length of VNTR and traits or expression. An alternative study in schizophrenia found an association based on the presence of different motif alleles rather than length of an intronic VNTR for the *CACNA1C* gene, suggesting that identifying VNTR alleles based on breakpoint clustering can miss important associations based on composition rather than length.

The accuracy of modern long-read sequencing and their assemblies enables the ability to study VNTR sequences according to their motif composition rather than breakpoints in sequence alignments. The motif composition of a sequence was introduced for deciphering the order of monomers that make up centromeric DNA (Dvorkina *et al*., 2020). This study extended wrap-around dynamic programming from single motifs (Fischetti *et al*., 1992; Loving *et al*., 2017) to multiple motifs in the StringDecomposer algorithm that takes as input a query sequence and a library of repeat motifs, and annotates the motif composition according to the optimal composition of repeat units or their subsequences that when concatenated together have the closest edit distance to the query.

The StringDecomposer method requires a database of known repeat motifs. Here we develop an automated approach for constructing a library of motifs from a reference set of haplotype-resolved assemblies. Naively, this list may be formed as the non-redundant set of repeated sequences identified by a repeat discovery tool such as Tandem Repeats Finder (Benson, 1999) ran on a reference set of VNTR sequences. However, this full list containing rare motifs can obscure the pattern of repetition. For example, the string ACGGTACGGTACCGTACGT may be decomposed into [ACGGT, ACGGT, ACCGT, ACGT], however (`ACGGT`)_4_ (4 repetitions) is more concise and has a low divergence from the original sequence. In a high level, we define the efficient motif set as a subset of original motifs in which rare motifs are replaced by more common ones such that divergence between VNTR sequences in the reference set and their decomposition by efficient motifs is bounded. For any new genomes, their annotations using efficient motifs are likely to remain within the divergence bound, given the assumption that the reference set is a large sample of the population diversity. To find such motif sets, we have developed a toolkit, <u>V</u>NTR <u>A</u>nnotation Using Efficient <u>Mo</u>tifs <u>S</u>et (vamos) that discovers motifs using a reference set of genomes, and finds efficient motif sets given a divergence parameter. Examples of VNTR decompositions using all observed motifs compared to efficient set of motifs annotations are shown in Fig 1; the *ACAN* intronic VNTR found to be associated with height(Beyter *et al*., 2021; Mukamel *et al*., 2021), and an intronic VNTR in WDR7 found to be a modifier of amyotrophic lateral sclerosis.(Course *et al*., 2020)

**Fig. 1.**
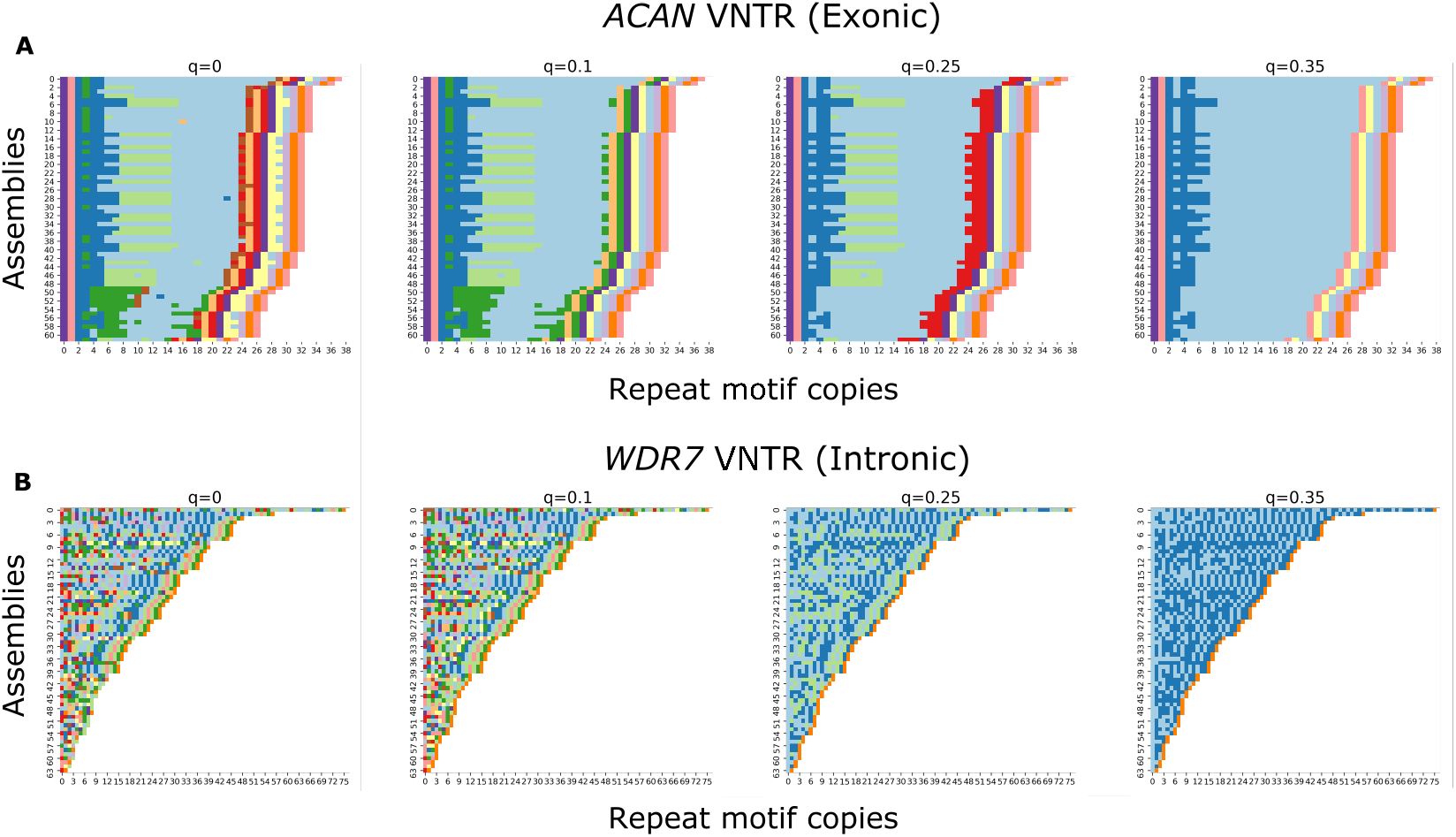
Visualization of allele structure change with four levels of divergence parameter Δ determined by parameter _*q*_. 64 HGSVC assemblies were annotated into a run-length encoding of efficient motifs selected at each level of Δ using StringDecomposer algorithm. Repeat motifs is color-coded and each individual allele is plotted as a series of colors (row). Repeat motif unit is shown on x-axis. The efficient motif set selected at *q* = 0 is basically the original motif set, since corresponding Δ = 0. With Δ growing larger, only truly representative motifs are selected and rare/contamination motifs possibly resulting from sequence error disappear and the allele’s domain structure starts to grow more clear. A. ACAN (chr15:88855422-88857301 on GRCh38) repeat alleles ranked by decreased length. B. WDR7 (chr18:57024494-57024955 on GRCh38) repeat alleles ranked by decreased length.

We generated set of efficient motifs for each VNTR locus under three levels of divergence parameter (Δ) from 64 haplotype-resolved assemblies sequenced with long reads (LRS) by the Human Genome Structural Variant Consortium.(Ebert *et al*., 2021, HGSVC,) As a proof of concept, we use our method to quantify diversity of VNTR sequences in the haplotype-resolved assemblies, and evaluate the application of the StringDecomposer method with efficient motif sets on long reads and simulated new genomes.

## 2 Materials and Methods

### 2.1 Generating a reference motif set

The efficient motif set is calculated on a panel of haplotype-resolved assemblies using sequences orthologous to VNTR loci in the reference (e.g. GRCh38). Given the location of a single VNTR in the reference, whole-genome alignments are used to determine the corresponding boundaries in the assemblies. The VNTR sequence in the reference and all orthologous VNTR sequences from assemblies are collectively referred to as a single VNTR locus. Each tandem repeat locus is annotated using Tandem Repeats Finder (Benson, 1999, TRF,) to determine an initial set of motifs observed for the locus across assemblies.

Each TRF annotation includes a repeat sequence and a consensus for the motif. The motif composition is additionally parsed from TRF annotations and maintained as a list of motifs that when concatenated form the originally annotated sequence. Due to changes in repeat pattern or other interruptions in repeat sequence, TRF may report multiple annotations for a single locus.

We employed a set of practical filters so that only one annotation is used per sequence, and for any annotation, redundant and partial motifs are excluded. First, annotations that are strictly homopolymer or di-nucleotide motifs are trivial to decompose and are removed. Next, annotations spanning less than 50% of the locus or having less than 80% base matches to the computed motif consensus are additionally excluded. Finally, redundant calls that are a multiplex of a smaller call are excluded by filtering calls where the length of the consensus motif is 1.5 times longer than the consensus motif of another annotation, and encompassed more than 80% the smaller motif. If multiple annotations passed these two filters for a sequence, the annotation where the least divergence between the repeat motifs and the consensus sequence measured as the standard deviation of the edit distances between the motifs and the consensus is selected as the representative annotation for the sequence. The annotation-level filtering is applied independently for each locus.

The full set of reference motifs for a locus are formed as the nonredundant union of individual sequence motifs from representative TRF annotations after applying motif-level filtering. We found that the first and last motifs to be partial or redundant. To account for this motifs on the VNTR boundaries that were longer than 2*/*3 or shorter than 1*/*3 of the motif consensus are eliminated. Since repetitive regions are highly variable across individuals, motifs from different genomes may be relate as cyclical permutations. We detect cyclical shifts by comparing motifs to an overall motif consensus derived from an abPOA multiple sequence alignment (MSA) (Gao *et al*., 2021) on the individual consensus motif sequences from filtered TRF annotations. Denote the overall MSA consensus as *a* and one of the consensus motif sequences as *b*. To avoid comparing all instances of cyclic shifts, the number of shifting bases *n* can be obtained by performing a local alignment between *a* and *b*·*b* (the tandem duplication of *b*). If the local alignment gives perfect prefix match between *a* and *b* · *b, n* is taken as the number of bases in the unmatched suffix of *b* + *b* and vice versa. When neither perfect head nor tail match is returned by the alignment, *n* is taken as the number of bases in the unmatched head of *b* · *b*. Every motif corresponding to *b* is shifted by *n* accordingly. This adjusted all motifs to the same frame as the overall MSA consensus. To further avoid alternative sequence decomposition in annotating new sequences by the motif set, we also split motifs that may be formed as an exact string decomposition of other shorter motifs discovered in that locus. Finally, for computational efficiency in downstream analysis, we excluded loci in centromeric DNA or that were longer than 10kb.

The result of the annotation and motif level filtration steps are to harmonize tandem repeat motif annotations that are sometimes inconsistent between samples. However, the filtration also has the effect that some of the original sequences defined by orthologous intervals to VNTR loci in the reference genome are not covered by tandem repeat annotations. We assume that the assemblies used to define loci is of sufficient size and quality that effective estimates of motif diversity remain despite any filtering. The problem for efficient motif set discovery is presented assuming the original VNTR sequences are fully annotated after filtering, or are modified to only represent subsequences of the original loci that remain annotated after filters are applied.

### 2.2 Efficient motif selection

The efficient motif selection applies independently across all VNTR loci. Here we consider efficient motif selection for one VNTR locus. Given a collection of orthologous VNTR sequences *V* = {*v*_1_, …, *v*_*J*_} from a reference set of *J* assemblies, the set of all distinctly observed motifs in *V* (after the filtering in section 2.1) is called original motif set, defined as Σ and Σ^*^ is the complete set of sequences that are concatenations of motifs from Σ. It follows that each *v*_*i*_ is a member of Σ^*^. An efficient motif set is a subset of Σ such that corresponding sequences may be composed from the efficient motif set that are similar to the original sequences. Specifically, given a divergence parameter Δ, an efficient motif set 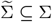 is a minimizer of the sum of 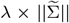 and 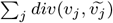 under the constraint that 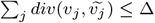, where *λ >* 0 is a hyper-parameter and 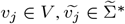, and *div* is the sequence divergence defined by edit distance.

Because Δ reflects total edit distance and not a percent identity, loci with longer or more divergent motifs require larger Δ for efficient motif set 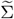 to be likely smaller than the original motif set Σ. Δ is designed to control the number of motifs that can be removed from original motif set Σ and set a reasonable removing cost per motif based on the motif divergence at each locus. Let *M* be the full list of all observed motifs in *V*. A user-specified global parameter *q* ∈ [0, 1] is used to calculate a locus-specific Δ by setting Δ = (*Q*(*q*) ×||*M* ||* *q*). *Q*(*q*), the *q*-quantile pairwise edit distance of all pairwise edit distances of motifs in *M* reflects the motif divergence at a locus, indicating the upper bound of the allowed removing cost per motif. The second term ||*M* || * *q* indicates the upper bound of number of motifs that can be removed.

The value of Δ grows with increasing *q*. In the limit, a single motif will be selected for 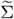 and the efficient motif set will lose the ability to represent variation in composition for a VNTR locus.

#### 2.2.1 Efficient motif selection problem

For one locus, let original motif set be Σ = {*m*_1_, *m*_2_, …, *m*_*p*_}, and counts that each motif appears in *V* be *O* = {*o*_1_, *o*_2_, …, *o*_*p*_}. Similar to the edit distance on sequences, the cost function 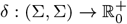 defines the cost of replacing a motif *m*_*i*_ by another *m*_*j*_, *δ*_*ij*_. The efficient motif set is formed by removing elements from Σ, and here consider replacing all instances of motif *m*_*i*_ by a motif *m*_*j*_ as how *m*_*i*_ is removed from Σ. We define the variable *x*_*ij*_ ∈ {0, 1} to indicate if *m*_*i*_ is replaced by *m*_*j*_. The objective of the motif selection is to minimize the sum of 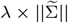 and total motifs replacement cost with the following three constraints. *λ >* 0 is a hyper parameter to balance the two terms. To roughly make the two terms at the same numeric level, we set *λ* to be equal to Δ in the actual computation.

1. Every *m*_*i*_ ∈ Σ is replaced by one and only one 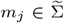 (possibly itself).
2. Motif *m*_*i*_ cannot be replaced by motif *m*_*j*_ if *o*_*i*_ *> o*_*j*_.
3. The total motifs replacement cost 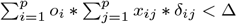.

Theorem 1 not only guarantees that if such efficient motif set 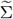 exists, the divergence between VNTR sequences {*v*_*j*_ |*j* ∈ *J*} and approximate sequences composed of only efficient motifs 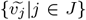 is bounded by Δ, but also gives a form of the composition of each 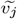. It is simply a concatenation of the counterpart efficient motif of each original motif, 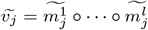, which is referred as *replacement annotation*.

##### Theorem 1.

*If an efficient motif set* 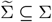 *exists, which minimizes the sum of* 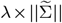 *and total motifs replacement cost* 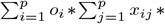 *δ*_*ij*_ *and satisfies the three requirements, then there exists* 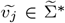 *such that* 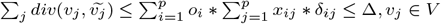.

Proof. Assume an efficient motif set 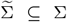 exists. Each VNTR sequence *v*_*j*_ can be represented as a sequence of original motifs 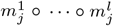, each 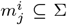. By substituting each 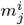 with its counterpart efficient motif 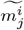, we can get 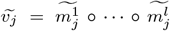. For each *v*_*j*_ and 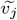, it is clear that 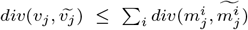, otherwise the combination of edit operations from individual motif alignments would infer a more parsimonious edit distance for *v*_*j*_ and 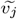. Therefore, 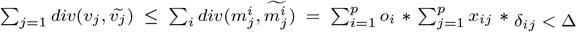.

#### 2.2.2 Integer linear programming formulation

Theorem 1 implies that it is possible to search for efficient motif set by bounding on the cost of replacing motifs independent of their context in *V*. We prove that the efficient motif set selection problem is a NP-hard problem when the cost function *div* is a general function not limited to edit distance in theorem 2 (Appendix). Fortunately, it can be formulated as an integer linear programming (ILP) problem (equation 2), which can be efficiently solved using the Google OR-tools (CP-SAT solver) (Google, 2019).

The problem may be formulated as a linear programming (LP) problem with an indicator function as objective (equation 1), which is not yet in ILP form due to the presence of indicator an function in the objective.

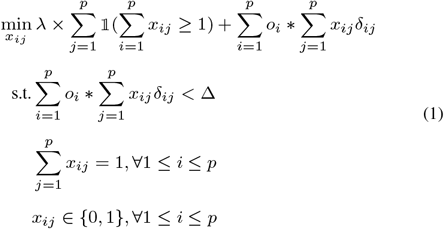

However, equation 1 can be transformed to an equivalent equation 2 by introducing additional variables 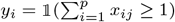. *L* and *Q* in equation 2 are the lower bound and upper bound of 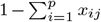 (In this case, *L* can be 1 − *p* and *Q* can be 1). Equation 2 is clearly in ILP form.

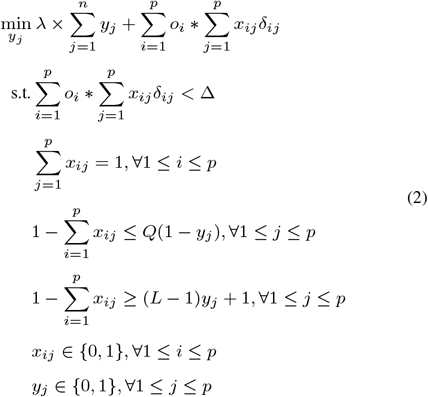

### 2.3 Annotating motif composition on sequence data

The vamos software adapts the StringDecomposer algorithm to annotate the motif composition of tandem repeat sequences for the use cases of alignments of haplotype-resolved assemblies and long (single-molecule sequencing) reads to a reference genome. As input vamos requires sequence alignments and a file with coordinates of reference VNTRs and a motif list for each VNTR. The tandem repeat sequences to annotate are extracted from the input alignments. When annotating assemblies, sequences are annotated directly as read from the alignments. When annotating read alignments, all reads covering a VNTR locus are first collected. Reads are partitioned by haplotype using phase tags if they have been phased using WhatsHap (Patterson *et al*., 2015) or HapCut2 (Edge *et al*., 2017)), or by a max-cut heuristic that initializes partitions with the two reads with the most disagreeing heterozygous SNVs and assigns remaining reads to a partition based on shared SNVs. The VNTR sequences from each of the reads are extracted based on alignments, and the StringDecomposer annotation is executed on the abPOA consensus (Gao *et al*., 2021) of the VNTR sequences in each partition. The output of a run is in Variant Call File (VCF) format (Danecek *et al*., 2011) with one variant entry per reference VNTR. The efficient motif list for a VNTR is stored in the `INFO` field along with the run-length encodings of each observed motif composition. The values of the genotype index the corresponding run-length encoding for each haplotype. In this manner, a combined-sample VCF maintains a record of distinctly observed alleles.

## 3 Results

### 3.1 Efficient motif discovery for a reference assembly panel

A set of 32 haplotype-resolved assemblies (64 haplotypes) was used as input for efficient motif set discovery (Ebert *et al*., 2021). The boundaries of 692,882 VNTR loci were obtained from the UCSC Genome Browser (Kent *et al*., 2002) for GRCh38. After filtering (2.1), there were 418700 ± 2714 annotated VNTR loci per genome. The union of annotations of all 64 assemblies yielded 467,104 annotated VNTR loci. On average, there were 4.8 ± 14.1 motifs per locus, of which 60,017 were trivial repeats with a single motif.

The filtered motif sets were used as input for efficient motif discovery. We generated efficient motif sets for *q* = 10%, 20%, and 30% for an increasing number of reference assemblies to examine the motif diversity at different levels of compression and population sampling. The genome-wide total motif count (motif set size) was used to gauge how the efficient motifs change according to the number of assemblies included in the discovery panel from eight to 64 assemblies. The size of the original motif set increases as genomes are added (Figure 2), as expected based on the inclusion of rare motifs. The efficient motif set size increases as more assemblies are included, with total diversity in this category showing asymptotic leveling at 64 genomes. After the inclusion of 40 assemblies, ∼ 98.36% efficient motifs are selected compared to the entire set when 64 assemblies are incorporated and *q* = 10% (Figure 2). On average, the efficient motif set size per VNTR locus ranges from 3.4 at *q* = 10% to 2.2 at *q* = 30% when 64 assemblies are included, while the original motif set size averages 5.4 per VNTR locus (Table 1). The mean compression ratio of motifs (efficient / original motif size) ranges from 0.81-0.66 (*q* = 10 − 30%) (Table 1, Figure 3).

**Table 1.** The mean efficient motif set size and compression ratio under three levels of Δ (10/20/30% quantile), when 64 haploid assemblies are incorporated.

| quantile | 10 | 15 | 20 |
| --- | --- | --- | --- |
| Mean efficient motif set size | 3.40 | 2.66 | 2.23 |
| Mean compression ratio | 0.81 | 0.72 | 0.66 |

**Fig. 2.**
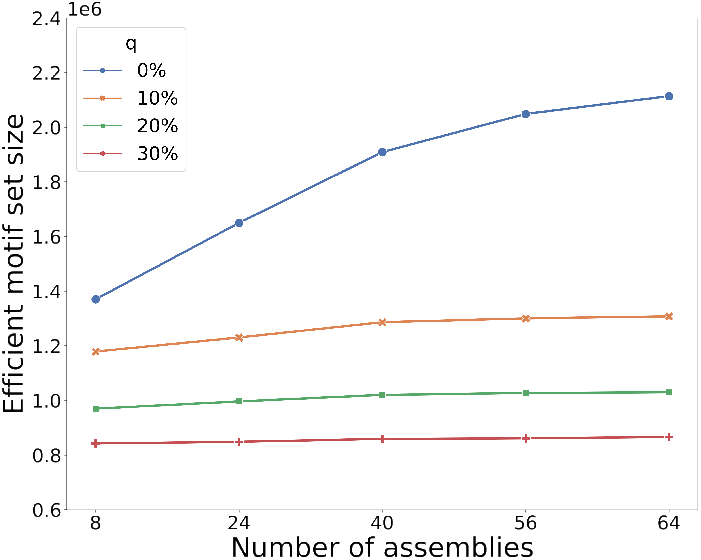
Diversity of efficient motif sets under different levels of compression as more assemblies are incorporated, measured by genome-wide size of efficient motif sets. Four levels of divergence parameter Δ are selected at *q* = 0*/*10*/*20*/*30%. The curve of *q* = 0% means no compression on the motif set and reflects the number of original motifs. As expected, the number of the original motifs increases as genomes are added, due to the inclusion of rare motifs. The efficient motif set size increases as more assemblies are included, with total diversity in this category showing asymptotic leveling at 64 genomes. As *q* grows larger, more compression is posed on the original motifs, resulting in less efficient motifs.

**Fig. 3.**
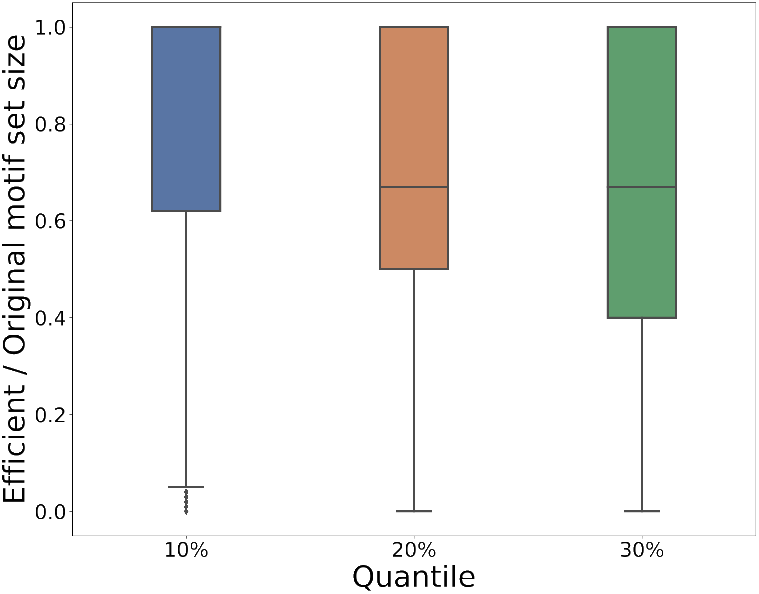
The compression ratio (efficient / original motif set size) under three levels of Δ (selected at *q* = 10*/*20*/*30% quantile), when 64 haploid assemblies are incorporated.

A simple approach to selecting an efficient motif set is to greedily select the most frequently occurring motifs at each locus. To evaluate the difference in annotations obtained when using efficient motifs selected by vamos and the simplistic approach, we annotated motif compositions of the 64 HGSVC haplotype-assemblies using the efficient motif sets and the same size of motif set selected by frequency. In total, 407,087 VNTR loci (excluding trivial repeats with a single motif) were annotated across 64 haplotype-assemblies, and were evaluated according to the edit distance between the nucleotide sequence translated from annotated motifs and the assembled sequence. We considered an annotation to be superior to another if its edit distance to the true VNTR sequence is less than 90% of the corresponding edit distance of the other sequence. On 6% (1,350,089) loci, the annotation using efficient motifs is superior than that using the greedy approach, which is roughly twice the number of loci where the greedy annotation is superior (Table 2). Compared with the greedy approach, efficient motifs have one advantage that it takes into account the cost of replacing each original motif, thus not only highly frequent motifs tend to be reserved but also less frequent ones with long length and dissimilar from other motifs. Suppose a locus has three motifs AAAG, AAAC, and TGTGACCTGCAC with counts 10, 3 and 1. The greedy approach picks top two frequent motifs `AAAG` and `AAAC`, whereas the efficient motifs are AAAG, TGTGACCTGCAC. In spite of the rarity of motif TGTGACCTGCAC, inclusion in the motif set will represent more diverse sequences that include this motif. The other advantage of efficient motifs is the automatic decision of the number of efficient motifs to select at each locus, which benefits from the ILP formulation and the association with a total replacement cost upper bound Δ which can adapt to the complexity of original motif set for each locus.

**Table 2.** Comparison of annotation using efficient motifs and baseline motifs. We define an annotation is significantly better than the other if the edit distance between the nucleotide sequence translated from the annotation in motif and the true VNTR sequence in assemblies is less than 90% of the other’s.

|  | # of loci |
| --- | --- |
| vamos annotation < greedy | 1,350,089 |
| vamos annotation < $0.9 \times$ greedy | 1,096,972 |
| greedy annotation < vamos | 744,547 |
| greedy annotation < $0.9 \times$ vamos | 569,509 |
| Total | 22,322,230 |

### 3.2 Allelic diversity in 64 haplotype-resolved *de novo* assemblies

We used vamos to study the allelic diversity of VNTR sequences in the 64 HGSVC haplotype-assemblies. Each assembly was annotated independently using the original motif set and efficient motif sets defined by *q* = 0.2. The average number of alleles per locus under the original motif set is 4.0, and under the efficient motif set is 3.1 (Table 3, Figure 4 A,B). The number of different alleles correlates with the average motif count using both the original (*r*^2^ = 0.31, *p <* 2.2 × 10^−16^) and efficient (*r*^2^ = 0.35, *p <* 2.2 × 10^−16^) motif sets. In contrast to annotations of alleles by either sequences or breakpoints, there are 3.5 alleles per locus when matching alleles by length, and 5.0 alleles per locus when comparing distinct sequences. Sequences were annotated using vamos v1.1.1.

**Table 3.** Average number of alleles per locus when genomes are annotated using the original/efficient set of motifs. Annotations are grouped into one allele if their edit distance with respect to the motif composition was 1-3 motifs.

| Motif set | Edit distance |  |  |  |
| --- | --- | --- | --- | --- |
|  | 0 | 1 | 2 | 3 |
| Original set | 4.0 | 2.7 | 2.1 | 1.8 |
| Efficient set ($q = 20\%$) | 3.1 | 2.3 | 1.8 | 1.6 |

**Fig. 4.**
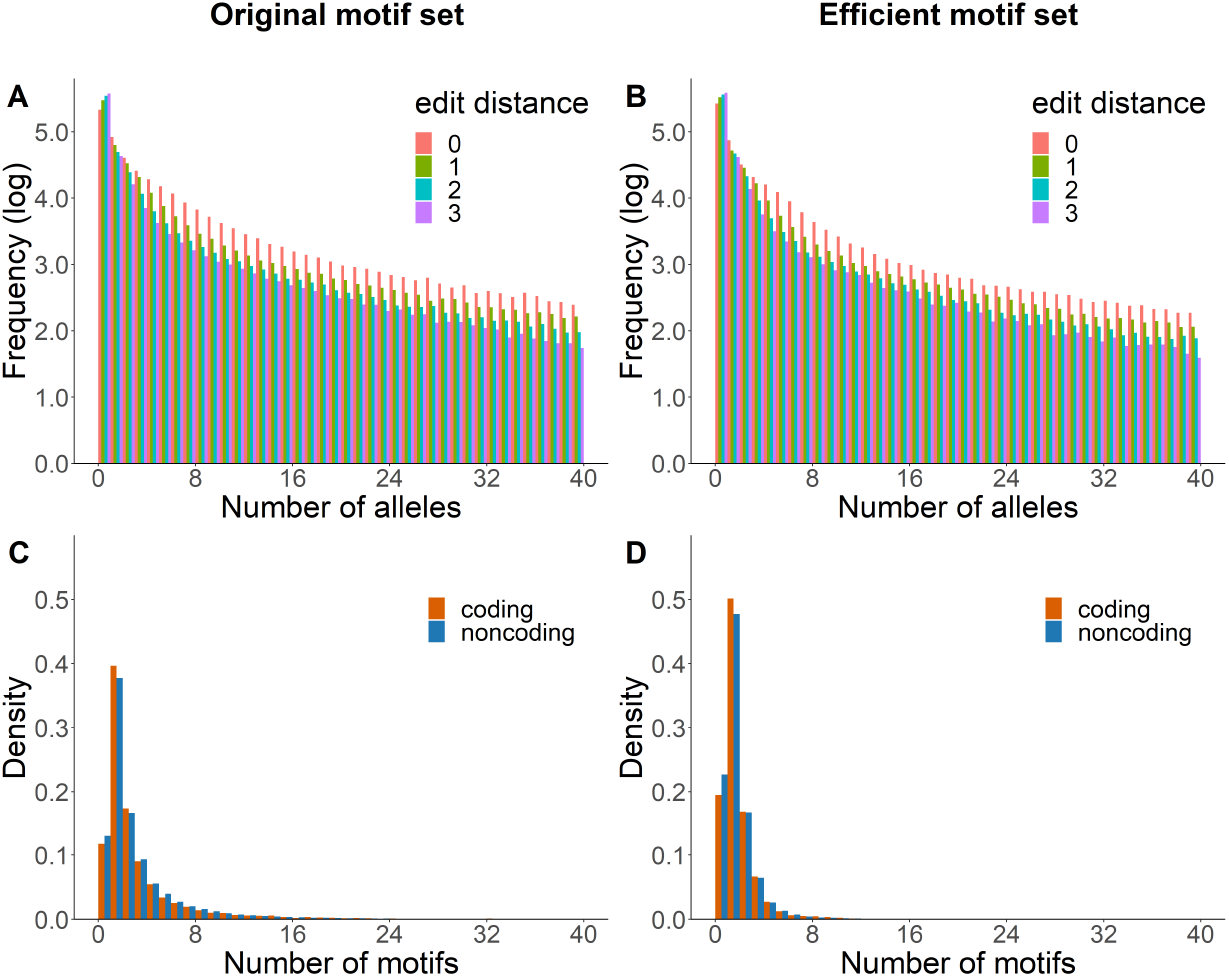
Distribution of number of VNTR alleles and unique motifs for 32 HGSVC diploid genomes. VNTR sequences are annotated by vamos contig using both efficient motifs (Δ selected at *q* = 20% quantile level) and original motifs. (A, B). Comparison of number of alleles when annotated by original motifs (A) and efficient motifs (B). Alleles are aggregated with maximum numbers of allowed edit distance of the annotation strings ranging from 0 to 3. (C, D). Comparison of number of unique motifs of coding/noncoding VNTRs in the original motifs set (C) and the efficient motif set (D).

To measure how different alleles are, we also grouped together annotations if their edit distance with respect to the motif composition was 1-3 motifs (Table 3, Figure 4 A,B). When considering annotations using the original motif set, there is a 47% reduction in the average number of alleles per locus when grouping alleles that differ by up to two motifs. Similarly, there is a 42% reduction in the average number of alleles in the efficient motif set under a similar grouping indicating that copy number variation instead of point mutation accounts for the majority of allelic variation, as expected by the slippage mutational mechanism of tandem repeat sequences.

We investigated the evolutionary constraint mutation in coding versus noncoding VNTR sequences. We found there was significantly lower diversity of alleles by length in coding sequences (*p <* 2.2 × 10^−16^, KS-test). However, there was no statistical difference in the motif diversity in coding versus noncoding VNTRs for the efficient motif sets (*p* = 0.07, KS-test, Figure 4 D), as expected with the decreased motif diversity, but also the original motifs (*p* = 0.07, KS-test, Figure 4 C). This could be due to a likely lower rate of mutation to create new motifs leading to a difference that is not detected in the sample size used, or a constraint on mutation in coding sequences.

### 3.3 Analysis of aligned long-reads

We analyzed long-read data of NA24385 sequenced at 10-30X for both HiFi and ONT platforms. Since analysis of assembly data gives the best overall performance by simulation, we compared annotations from reads data to those from the HGSVC NA24385 assembly (Figure 5). Poor coverage usually leads to unresolved phasing. Consequently, increasing sequencing depth from 10X to 20X greatly decreases the number of un-annotated loci for both platforms, yet such benefit is not as significant when sequencing depth is further increased from 20X to 30X. Under a sequencing depth of 30X, the percentage of VNTR loci with over 80% of the covered reads successfully phased is 76% and 70% for the HiFi and ONT data, respectively. However, although both platforms have over 98% of the VNTR loci covered at 10X or above, this number drops to 92% for the ONT data but 83% for the HiFi data after phasing, showing the advantage of longer reads for phasing. The HiFi data generally has more loci un-annotated than the ONT data, though for annotated loci the HiFi data produces better agreement to the assembly annotations. Furthermore, the unannotated loci include regions that may be autozygous and could be analyzed as a single homozygous annotation.

**Fig. 5.**
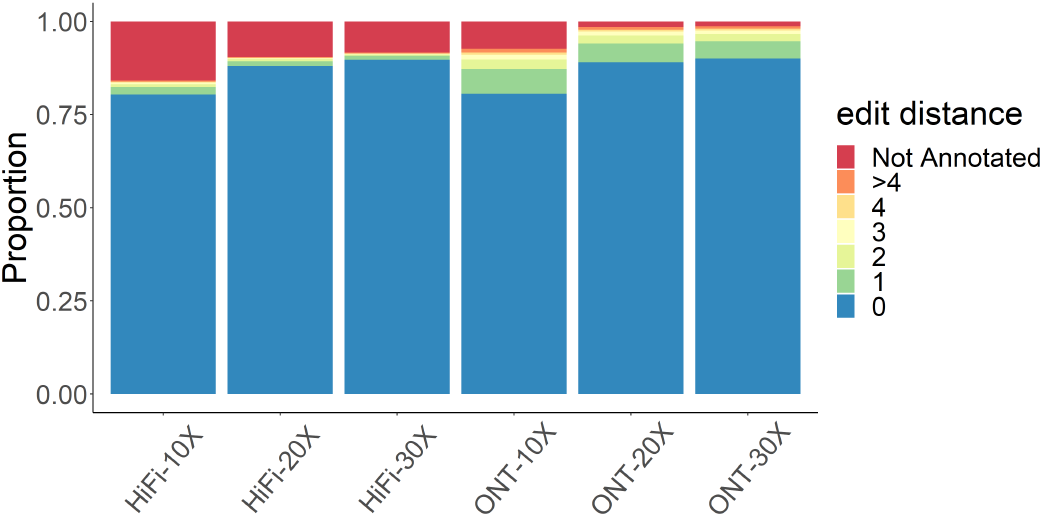
Analysis of raw sequencing reads of NA24385. Raw sequencing reads of NA24385 were annotated by vamos –read. Edit distance was calculated by aligning annotation strings to results from the HGSVC NA24385 assembly for each VNTR locus.

### 3.4 Annotating diverged populations

We used simulation analysis to evaluate annotations on alleles that have differences from the reference assemblies. VNTR sequences were simulated based on 500 randomly selected loci described in Section (2.1) and the corresponding replacement annotations were compared with the StringDecomposer annotations. Motif sets with a single motif were excluded from simulation since variations introduced by simulation may largely be concealed by the motif homogeneity in such cases. From each of the 500 selected motif sets, VNTR sequences were generated by sampling and concatenating 50-100 motifs according to their underlying frequencies in the reference assemblies.

Population diversity where sequences have motifs not reflected in the original or efficient motif sets was simulated by adding random mutations into the simulated VNTR sequences. Specifically, four settings for mutation rate: a 1% and 2% mutation rate sampled uniformly from single base substitution, deletion, or duplication combined with an optional 1% rate of insertions 2-4 bases.

The annotations were assessed using simulated assemblies (error-free), as well as 30-fold coverage of reads from HiFi using pbsim (Ono *et al*., 2013) with average 98.1% accuracy and ONT using alchemy2 (distributed with lra and averaging 88.8% accuracy). Similar to the analysis of aligned reads, the accuracy of the annotation is impaired by sequencing error (Figure 6). Under all conditions, the relative difference (motif edit distance / number of motifs in replacement annotation) is typically below 5%. The instances where there was a high divergence between the simulated motifs and the motif decomposition were typically due to long motifs in the original motif set being decomposed by shorter motifs in the efficient motif set.

**Fig. 6.**
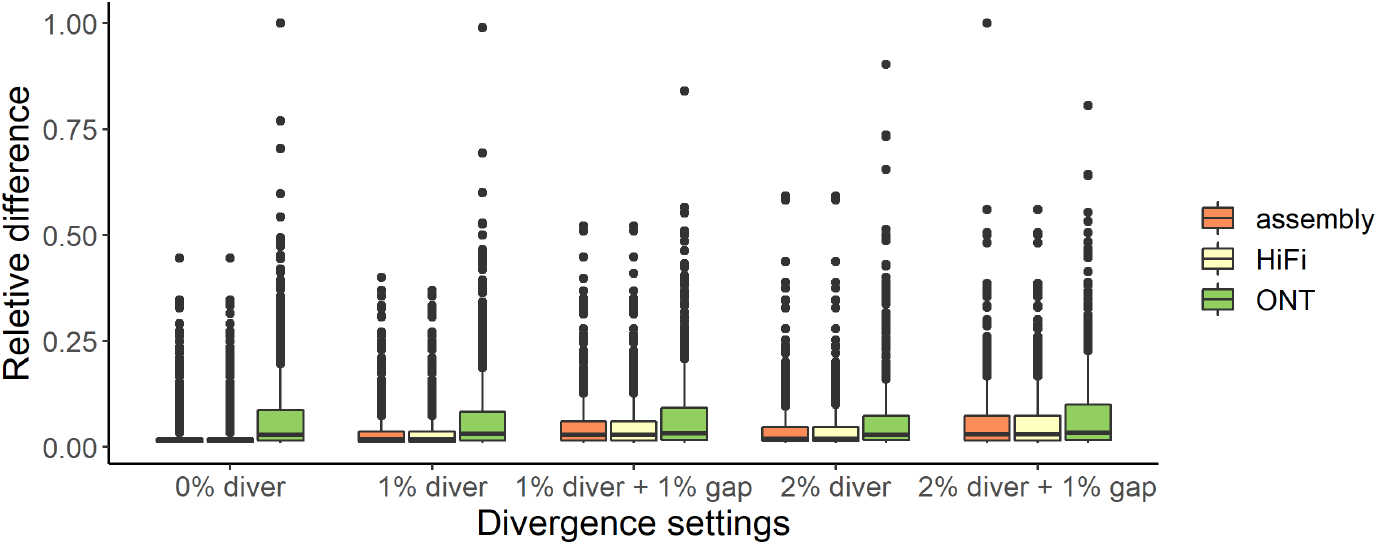
Relative difference between replacement annotation and StringDecomposer annotation on simulated assemblies, HiFi and ONT raw reads using efficient motif set (selected at *q* = 20% quantile) under five divergence settings. Relative difference is defined as the ratio of edit distance with respect to motif composition between two annotations to the length of replacement annotation.

## 4 Discussion

New long-read sequencing and assembly efforts are generating large databases of structural variation enriched at variable number of tandem repeats. These variants tend to be multi-allelic, however current catalogs of variation merge multiple different variant calls into a single representative call that limits the ability to associate different alleles with traits. One solution to this is to consider each unique VNTR sequence as an allele. Under this estimate, there are an average of 5 alleles per locus in VNTRs among the set of loci examined in this study. Here, we show there is a 50% reduction in annotated allele diversity when small-scale variation is abstracted by the efficient motif set annotation. This difference is subtle, but as the number of genome assemblies increases, the benefit of an abstract encoding of VNTR alleles will also grow.

The set of regions that vamos calls depends on the initial preprocessing of assemblies. The number of loci annotated in each genome depends on the overall quality of the assembly and the contiguity of the whole-genome alignments between the assembly and the reference. Future annotations based on the CHM13 T2T assembly (Nurk *et al*., 2022) will likely increase the number of loci included in the efficient motif database, and higher quality assemblies will expand the regions where an annotation database may be constructed, such as VNTRs in segmental duplications.

The appropriate parameters for selecting an efficient motif set will be specific to the context which the analysis is performed. Generally, for the reference assemblies that were considered in this study, values of *q* ≤ 20% preserved common motifs and excluded rare ones. Depending on the aims of the study, a higher value of *q* will enable characterization of domain structure or other higher-order patterns in repetitive DNA, and low, or complete motif sets can be used to study patterns of rare variation and mutation rates.

The runtime of ILP solver is highly variable across loci. It takes a few minutes to run for loci with up to hundreds of motifs, but for loci with thousands of motifs, it can take hours. However, each locus may be calculated in parallel, and it only needs to run once on a set of reference genomes. The vamos software is distributed with efficient motif sets computed for *q* = 10, 20 and 30%, along with the original motif set for the 64 haplotype-assemblies, and may be updated as additional human genomes are assembled.

## 5 Appendix

### Theorem 2.

*The efficient motif set selection problem is NP-hard*.

Proof. To prove the efficient motif set selection problem is NP-hard, we can prove a special case of the problem is NP-hard. When *λ* is set to ∞, the minimization objective turns into simply minimizing the size of the efficient motif set. When all motif counts *o*_*i*_ are the same, the second requirement in 2.2.1 will always be satisfied.

The decision version of simplified efficient motif set selection problem can be formulated in the following way: given a *k*, can we find a subset 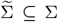 with 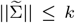, such that the total replacing cost 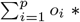 *x*_*ij*_ * *c*_*ij*_ < Δ. The decision version of the set cover problem can be formulated as the following: given a *k*, a set of elements {*e*_1_, *e*_2_, …, *e*_*p*_} and a collection of *p* subsets {*S*_1_, *S*_2_, …, *S*_*p*_} of elements, whose union equals the universe, can we find a sub-collection of *p* sets whose union equals the universe and the size of the sub-collection is less than or equal to *k*.

To prove the NP-completeness of the decision version of simplified efficient motif set selection problem, first, we show the problem belongs to *NP*.

Suppose we have a solution of variables {*x*_*ij*_}_*i,j*=1,…,*p*_, we can check if 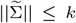 and the total cost 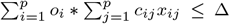 in polynomial time.

Second, we prove the problem is NP-hard by reducing the decision version of the special set cover problem with equal size of elements and sets to it.

Let a set cover problem instance be a set of *p* elements {*e*_1_, *e*_2_, …, *e*_*p*_} and a collection of *p* sets of elements {*S*_1_, *S*_2_, …, *S*_*p*_} whose union equals the universe of elements. For any set cover problem instance, we can create an instance of efficient motifs selection problem: define a set of *p* motifs as {*m*_1_, *m*_2_, …, *m*_*p*_} and associated counts *o*_*i*_ = 1 for ∀*i* = 1, …, *p*. Set Δ = 0.5 and *c*_*ij*_ = 𝟙(*e*_*i*_ ∉ *S*_*j*_).

Suppose {*x*_*ij*_}_*i,j*=1,…,*p*_ is a solution of the constructed instance of the efficient motifs selection problem. We can prove a sub-collection of sets which is defined as 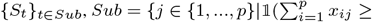 1)} is a valid solution of the set cover instance.

- Since 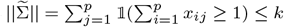, thus ||*Sub*|| ≤ *k*.
- Since each motif *m*_*i*_ must have a replacement, there exists some *j* such that *x*_*ij*_ = 1. Since 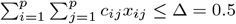 and *c*_*ij*_ = 𝟙(*e*_*i*_ ∉ *S*_*j*_), then *x*_*ij*_ = 1 ⇒ *c*_*ij*_ = 0 ⇒ *e*_*i*_ ∈ *S*_*j*_. Additionally by the definition of *Sub*, such *S*_*j*_ ∈ {*S*_*t*_}_*t*∈*Sub*_. Therefore, for each *e*_*i*_, there exists *S*_*j*_ ∈ {*S*_*t*_}_*t*∈*Sub*_ such that *e*_*i*_ ∈ *S*_*j*_.

Next, suppose a sub-collection {*S*_*t*_}_*t*∈*Sub*_, *Sub* ⊆ {1, …, *p*} is a solution of the set cover problem instance. We define {*x*_*ij*_}_*i,j*=1,…,*p*_ (variables indicating if motif *m*_*i*_ is replaced by *m*_*j*_) as follows:

- If *j* ∉ *Sub, x*_*ij*_ = 0.
- If *j* ∈ *Sub* and *e*_*i*_ ∉ *S*_*j*_, *x*_*ij*_ = 0.
- If *j* ∈ *Sub, e*_*i*_ ∈ *S*_*j*_, and *S*_*j*_ is the only set in {*S*_*t*_}_*t*∈*Sub*_ that covers *e*_*i*_, then *x*_*ij*_ = 1.
- If *j* ∈ *Sub, e*_*i*_ ∈ *S*_*j*_, and there are more than one set in {*S*_*t*_}_*t*∈*Sub*_ that covers *e*_*i*_, suppose 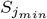 has the minimum index among these sets, set 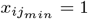 for any other *S*_*j′*_, set *x*_*ij′*_ = 0.

We next prove the defined {*x*_*ij*_}_*i,j*=1,…,*p*_ is a valid solution of the constructed instance of the efficient motif selection problem.

- 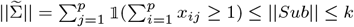.
- For each *e*_*i*_, there must exist some *j* such that *e*_*i*_ ∈ *S*_*j*_. So there must exist some *j*^′^ (not necessarily *j*) such that *x*_*ij′*_ = 1. And according to the definition, there is only one *j*^′^ such that *x*_*ij′*_ = 1.
- For every *i, j*, since 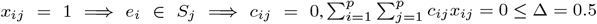.

To sum up, we prove the decision version of simplified efficient motif selection problem is NP-complete, thus the optimization version of the problem is NP-hard. Therefore, efficient motif selection problem is NP-hard.

## Acknowledgements

We thank Andrey Bzikadze and Pavel Pevzner for their useful comments on StringDecomposer analysis.

## Data Availability

- 64 haplotype-resolved assemblies produced by the Human Genome Structural Variation Consortium (Ebert *et al*., 2021) can be downloaded from http://ftp.1000genomes.ebi.ac.uk/vol1/ftp/data_collections/HGSVC2/release/v1.0/assemblies/.
- The NA24385 data is available from NCBI project PRJNA586863.
- Motif set files and combined vcf of VNTR annotations are available through zenodo at doi:10.5281/zenodo.7158427.

